# Human gray matter microstructure mapped using Neurite Exchange Imaging (NEXI) on a clinical scanner

**DOI:** 10.1101/2024.10.04.616639

**Authors:** Quentin Uhl, Tommaso Pavan, Thorsten Feiweier, Gian Franco Piredda, Ileana Jelescu

## Abstract

Biophysical models of diffusion in gray matter (GM) can provide unique information about microstructure of the human brain, in health and disease. Therefore, their compatibility with clinical settings is key. Neurite Exchange Imaging (NEXI) is a two-compartment model of GM microstructure that accounts for inter-compartment exchange, whose parameter estimation requires multi-shell multi-diffusion time data. In this work, we report the first estimates of NEXI in human cortex obtained on a clinical MRI scanner. To do that, we establish an acquisition protocol and fitting routine compatible with clinical scanners.

The model signal equation can be expressed either in the narrow-pulse approximation, NEXI_NPA_, or accounting for the actual width of the diffusion gradient pulses, NEXI_WP_. While NEXI_NPA_ enables a faster analytical fit and is a valid approximation for data acquired on high-performance gradient systems (preclinical and Connectom scanners), on which NEXI was first implemented, NEXI_WP_ has significant relevance for data acquired on clinical scanners with longer gradient pulses. We establish that, in the context of broad pulses, NEXI_WP_ estimates were more comparable to previous literature values.

Furthermore, we evaluate the repeatability of NEXI estimates in the human cortex on a clinical MRI scanner and show intra-subject variability to be lower than inter-subject variability, which is promising for characterizing healthy and patient cohorts.

Finally, we analyze the relationship of NEXI parameters on the cortical surface to the Myelin Water Fraction (MWF), estimated using an established multicomponent T_2_ relaxation technique. Indeed, although it is present in small quantities in the cortex, myelin can be expected to decrease permeability. We confirm a strong correlation between the exchange time (*t*_ex_) estimates and the MWF, although the spatial correspondence between the two is brain-region specific and other drivers of *t*_ex_ than myelin density are likely at play.

## 1. Introduction

Characterizing the brain’s microstructure has a potentially far-reaching impact on the clinical management of neurodegenerative diseases. Diffusion MRI models offer a unique approach to probing the brain’s tissue architecture, revealing subtle changes that often go undetected with conventional imaging techniques (Jelescu and Fieremans, 2023; Jones, 2010; Pavan et al., 2024). These models can discern alterations in neuronal and glial cells, myelination, and extracellular matrix properties (Alves et al., 2022; Hutchinson et al., 2018; Jelescu et al., 2016b; Vestergaard-Poulsen et al., 2007; Wu et al., 2024), which are critical for understanding the progression of diseases such as Alzheimer’s (Dong et al., 2020; Falangola et al., 2013; Gong et al., 2013; Pavan et al., 2023; Tristão Pereira et al., 2021), Parkinson’s (Atkinson-Clement et al., 2017; Sejnoha Minsterova et al., 2020; Zhan et al., 2012), multiple sclerosis (MS) (Liao et al., 2024; Martínez-Heras et al., 2020), first-episode psychosis (Rae et al., 2017) and schizophrenia (Kraguljac et al., 2019; Pavan et al., 2024). The ability of diffusion MRI to reveal and characterize microstructural changes early and accurately could lead to more timely interventions, personalized treatment plans, and better overall patient outcomes.

For white matter, most of the biophysical models of diffusion are variants of a well-accepted general model, the Standard Model of white matter (Novikov et al., 2019).

For gray matter, new models have been recently proposed, complementing the Standard Model either by adding the contribution of cell bodies modeled as spheres (SANDI) (Palombo et al., 2020), of exchange between neurites and extracellular water (NEXI, SMEX) (Jelescu et al., 2022; Olesen et al., 2022), or both (SANDIX) (Olesen et al., 2022). The feasibility of GM models has so far been demonstrated mainly on MRI systems benefitting from high-performance gradients, such as preclinical scanners or human Connectom scanners with maximum gradient amplitudes of ≥300 mT/m (Palombo et al., 2020; Uhl et al., 2024b).

Experimental observations of decreasing MR signal with increasing diffusion times in the gray matter suggest exchange as the dominant contributor over restricted diffusion within soma (Jelescu et al., 2022; Olesen et al., 2022). This has implications for the applicability of SANDI, which, by neglecting exchange, is best suited for short diffusion times where the assumption of impermeable compartments remains valid. Moreover, the effects of soma restriction become particularly prominent at high b-values, highlighting the necessity of acquiring high b-value data to effectively probe GM features. Initial attempts to implement SANDI in a clinical setting have been made (Schiavi et al., 2023), demonstrating the growing interest in translating these advanced models to more accessible platforms.

As a parallel attempt to model gray matter, NEXI does not model soma but rather accounts for the inter-compartment exchange, therefore overcoming the need for short diffusion times. However, this comes at the cost of requiring the acquisition of multiple diffusion times. This model characterizes neurites as a collection of randomly oriented sticks, occupying a relative signal fraction *f*. The intra-neurite diffusion is modeled as unidirectional with a diffusivity *D*_*i*_, while the extra-neurite compartment is modeled as Gaussian isotropic with diffusivity *D*_*e*_. Water exchange between these two compartments occurs with a characteristic time *t*_ex_.

The NEXI signal equations can be derived under two different assumptions: the Narrow Pulse Approximation (NEXI_NPA_) (Jelescu et al., 2022) and the actual Wide Pulses (NEXI_WP_, implemented in (Olesen et al., 2022)). NEXI_NPA_ is a valid approximation for data acquired on preclinical scanners (Jelescu et al., 2022; Olesen et al., 2022) and on human Connectom scanners (Lee et al., 2022; Uhl et al., 2024b), with gradient pulse duration (δ) values of, e.g., 4 and 9 ms, respectively. However, this assumption is challenged for clinical protocols where δ typically needs to be much longer to compensate for lower gradient amplitude and potentially different stimulation limits.

Here, we propose a clinically feasible NEXI protocol and evaluate its performance in characterizing human cortical microstructure non-invasively. First, we evaluate the impact of NEXI_NPA_ vs NEXI_WP_ on microstructure estimates. Second, similarly to our approach on a Connectom system (Uhl et al., 2024b), we evaluate the repeatability of NEXI estimates on a clinical scanner by comparing intra-subject and inter-subject variability using scan-rescan of a cohort of healthy volunteers. Third, we explore for the first time correlations between NEXI-derived cortical microstructure features and complementary techniques such as Myelin Water Imaging. We hypothesized that one of the main drivers of cell membrane permeability (and thus exchange time *t*_*ex*_) would be myelin content, which is also found in the cortex (Nieuwenhuys, 2013; Palomero-Gallagher and Zilles, 2019), and is expected to reduce permeability. The inter-compartment exchange time could thus serve as an innovative proxy for assessing the density of myelin in gray matter. To robustly quantify myelin using clinical MRI, we used the T_2_-based approach of (Piredda et al., 2021b) to estimate the Myelin Water Fraction (MWF).

## 2. Methods

### 2.1 Theory: NEXI model variants

In both NEXI_NPA_ and NEXI_WP_, the signal results from the integration of a kernel around several directions.

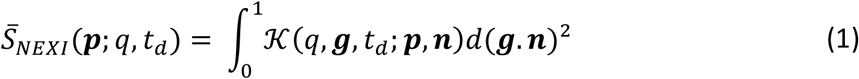

where ***p*** = [*t*_*ex*_, *D*_*i*_, *D*_*e*_, *f*] are the microstructure parameters to fit, **n** are the neurite orientations, *q* is the wave vector along direction **g** (Jelescu et al., 2022; Olesen et al., 2022).

Each method has its own way of deriving the kernel 𝒦. NEXI_NPA_, employing the Kärger model approximation (Kärger, 1985), simplifies the complex exchange dynamics by treating diffusion encoding pulses in the Pulsed Gradient Spin Echo (PGSE) design as instantaneous Dirac pulses, separated by a duration Δ. This approximation facilitates a more straightforward analytical solution but at the cost of accuracy. A correction term is introduced, however, by considering the diffusion time to be t_d_ = Δ-δ/3, instead of Δ (Moutal et al., 2018).

On the other hand, NEXI_WP_ adopts a more theoretically comprehensive stance by numerically integrating the system over the duration of the gradient pulses, acknowledging the real pulse width. Its signal was calculated using the Initial Value Problem (IVP) solver (*solve_ivp* from scipy.integrate (Virtanen et al., 2020)). While this approach enhances model fidelity to actual experimental conditions, it demands higher computational resources and faces challenges in numerical stability and precision. An efficient way to speed up the fit is to solve the equation on the segment [δ, Δ] whose solution is :

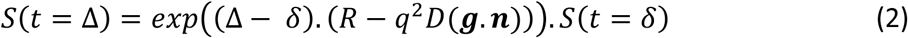

where *R* and *D* are respectively the rate and diffusivity matrices of the generalized rate equation (Ning et al., 2018), as well as to jointly calculate the signal and its analytical Jacobian and to use the tensor formalism. These formulae are provided in our code repository (https://github.com/Mic-map/graymatter_swissknife).

### 2.2 Experimental

#### 2.2.1 Participants

The study was approved by the ethics committee of the canton of Vaud, Switzerland (CER-VD). Written informed consent was obtained from all participants. Data were acquired in eleven healthy adults (6M, 26.9±1.3 years old). Each participant was scanned twice (delay between scans: 65 ± 29 days).

#### 2.2.2 Data acquisition

All data were acquired on a 3T MRI system with 80 mT/m gradients (MAGNETOM Prisma, Siemens Healthineers AG, Forchheim, Germany). An anatomical reference was acquired using an MP-RAGE sequence (1-mm isotropic resolution, FOV = 224 × 240 mm^2^, 256 slices, TI/TR = 900/1760 ms). Diffusion-weighted images were acquired using a PGSE Echo-Planar Imaging (PGSE EPI) research sequence with b-values = 1.00 and 2.00 ms/µm^2^ at diffusion times Δ = 28.3 and 36.0 ms, b-values = 1.00, 2.00, 3.20, 4.44 ms/µm^2^ at Δ = 45.0 ms, and b-values = 1.00, 2.00, 3.20 and 5.00 ms/µm^2^ at Δ = 55.0 and 65.0ms, 20 directions per shell, in addition to one b = 0 ms/µm^2^ images per Δ and one b = 0 ms/µm^2^ with reversed EPI phase encode direction for susceptibility distortion correction. Other parameters were fixed: δ = 16.5 ms, TE/TR = 100 ms/5 s, FOV = 256×256 mm^2^, matrix: 128×128, 64 slices, 2-mm isotropic resolution, partial Fourier = 6/8, GRAPPA = 2, SMS = 2. The total scan time for dMRI was 27 min. Multi-echo T_2_ data were collected using a 3D multi-echo accelerated gradient and spin echo (GRASE) research sequence (Piredda et al., 2021a) with voxel-size=1.8×1.8×1.8mm^3^; ΔTE/N-echoes=10.94ms/32; TR=5s, matrix-size=112×128×76.

#### 2.2.3 Diffusion data preprocessing

While each diffusion time was acquired in a separate acquisition run, all multi-shell multi-diffusion time data (N = 325 volumes) were pooled together for pre-processing. Pre-processing included Marchenko-Pastur principal component analysis (MP-PCA) magnitude denoising (Veraart et al., 2016), Gibbs ringing correction (Kellner et al., 2016), distortion and eddy current correction (Andersson and Sotiropoulos, 2016). A separate MP-PCA denoising of *b* = 0 and *b* = 1 ms/µm^2^ images (N = 112 volumes) was used to extract an unbiased noisemap, σ, from high SNR data, to be used in the Rician mean correction. For NEXI estimation, data were averaged over directions (powder-average, using the arithmetic mean) and normalized by the mean value of the *b* = 0 volumes.

#### 2.2.4 Cortical thickness and ROI parcellation

Cortical thickness was extracted from anatomical MPRAGE images using FastSurfer (Henschel et al., 2020). Grey matter region of interests (ROIs) from the Desikan-Killiany-Tourville (DKT) atlas (Klein and Tourville, 2012) were segmented on the anatomical MPRAGE image using FastSurfer and transformed into diffusion native space using linear registration of distortion-corrected b = 0 images to MPRAGE images. The cortical ribbon was segmented by merging the gray matter ROIs obtained with the DKT atlas. Mean values of NEXI parameters, MWF and cortical thickness were calculated for each ROI. The spatial distribution of these measures averaged across subjects was examined using inflated brain surfaces obtained from Connectome Workbench (Marcus et al., 2011).

### 2.3 NEXI_NPA_ and NEXI_WP_ parameter estimation and comparison

The expectation value of the Rician noise floor is incorporated directly into the model fit, as described in (Uhl et al., 2024b). The Rician floor markedly affects low-SNR images, i.e. high b-value images in our case. The Rician scale σ (which determines the Rician floor as 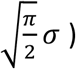 was fixed to the noise standard deviation estimated using MP-PCA during preprocessing.

The two NEXI model variants were fit to the experimental data by Nonlinear Least Squares (NLS) using the L-BFGS-B algorithm and *minimize* function (from *scipy*.*optimize* (Virtanen et al., 2020)) with a tolerance of 1e-14. The bounds specified for the optimization were [1 - 150] ms for *t*_ex_, [0.1 - 3.5] µm^2^/ms for the two diffusivities and [0.1 - 0.9] the fraction *f*. The initialization had the constraint that *D*_i_>*D*_e_ (Dhital et al., 2019a; Howard et al., 2022a; Kunz et al., 2018).

An initial grid search was applied before the NLS to find an optimal starting point. The parametric maps of the two variants (NEXI_NPA_ and NEXI_WP_) are compared on the inflated brain surface, as well as the averages of the parameter values in each ROI.

To compare the models’ goodness-of-fit to our experimental data, we used the corrected Akaike Information Criterion (AICc) (Akaike, 1973). A lower AICc suggests a better fit.

### 2.3 Performance Evaluation

#### 2.3.1 Noise propagation using synthetic signals generated from experimental distributions

To assess the propagation of noise in our estimates and how the NEXI_NPA_ model handled signals generated using wide gradient pulses, we used synthetic ground truth signals drawn from the parameter distributions obtained throughout the cortex with NEXI_WP_. We selected 10,000 voxels whose NEXI_WP_ estimates converged without hitting any bound, and whose diffusivities respected *D*_*i*_>*D*_*e*_. We then added Rician noise, with 20 repetitions to mimic the number of directions in the acquisition protocol, averaged them as done for the powder-average and estimated the NEXI_NPA_ and NEXI_WP_ parameter values using the same fitting procedure as for experimental data.

#### 2.3.3 Repeatability and brain region-specific patterns

Intra-subject vs inter-subject variability was assessed on average GM mean ROI estimates obtained by the NEXI variants. The different correlation coefficients between estimates from subjects and between sessions of the same subjects are also examined.

### 2.4 Myelin comparison

#### 2.4.1 Myelin Water Fraction estimation

MWF was estimated using the multicomponent T_2_ toolbox (https://github.com/ejcanalesr/multicomponent-T2-toolbox). Recent benchmark of non-parametric T_2_ relaxometry methods (Canales-Rodríguez et al., 2021) recommended the use of either χ^2^-I or L-curve-I to fit the T_2_ distribution, depending on the noise level. A comparison of the sharpness of the cut-off between the T_2_ distribution lobes with each technique led us to choose χ^2^-I (see Supplementary Fig. S6 for an example). MWF was therefore estimated using the χ^2^-I non-parametric T_2_ relaxometry method for myelin water quantification (Canales-Rodríguez et al., 2021).

#### 2.4.2 Relationships between MWF and NEXI parameters

We calculated the correlation between NEXI_NPA_ or NEXI_WP_ parameter estimates and MWF across ROIs, averaged over subjects. The cortical surface maps of *t*_*ex*_ and MWF are visually compared.

### 2.5 Relationships between cortical thickness and NEXI parameters

NEXI parameters estimates measured in a GM voxel might be influenced by Partial Volume Effects (PVE) with neighboring WM or CSF. As areas of thinner cortical thickness may be more affected by PVE due to the relatively large DWI voxel size (2mm), we therefore also evaluated potential correlations between the cortical thickness and the NEXI parameter estimates across ROIs.

## 3. Results

### 3.1 NEXI parameters in the human cortical ribbon

We report the first estimates of gray matter microstructure in the human brain obtained using NEXI and a clinical scanner (Figure 1). The two implementations, NEXI_NPA_ and NEXI_WP_, are compared. As anticipated, the observable patterns across brain regions are identical between the two implementations, with the only significant difference being a slight shift in the scales – i.e. somewhat lower estimates of *t*_ex_, *f, D*_i_ and *D*_e_ for NEXI_WP_ vs NEXI_NPA_. Both implementations however yield a *t*_*ex*_ between 30 and 60 ms in the cortex, which is on the order of the previous estimate of 42 ms on the Connectom scanner (Uhl et al., 2024b).

**Fig. 1.**
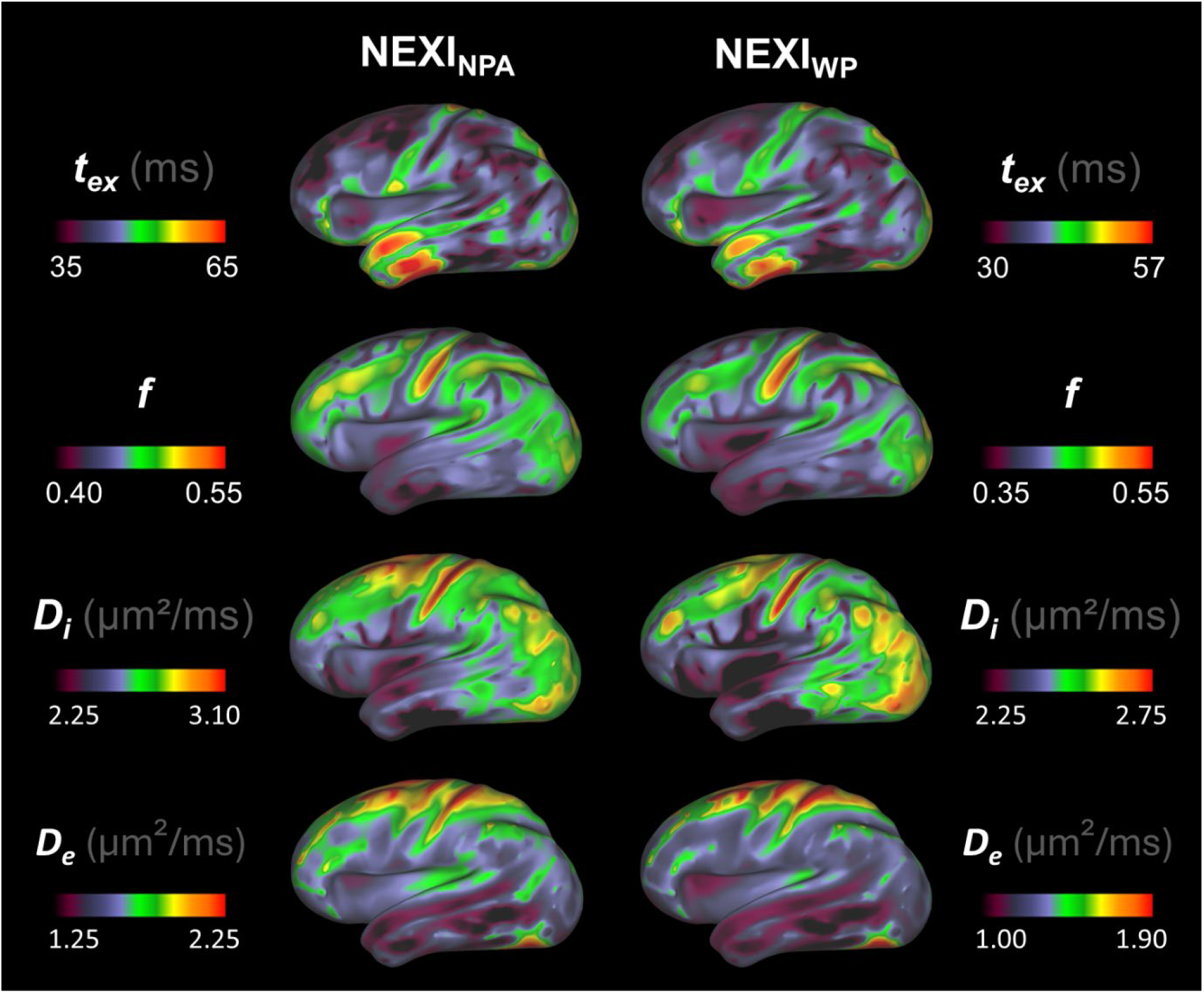
Projection onto cortical surface of NEXI_NPA_ and NEXI_WP_ maps averaged across subjects and sessions. While the overall patterns remained consistent between models, there is a noticeable shift in scale from one model to the other. As compared to other brain areas, higher values of *f, D*_*i*_, and *D*_*e*_ are recorded around the central sulcus, contrasting with lower values in the anterior temporal lobe.

The brain surface maps consolidate some patterns described in (Uhl et al., 2024b), notably along the central sulcus and the anterior temporal lobe. We further observe a pattern of longer *t*_*ex*_ in the temporal lobe, reaching values exceeding 50 ms, paired with a lower *D*_*i*_ estimate.

There are some notable differences with initial estimates on the Connectom scanner (Uhl et al., 2024b). For example, higher *D*_*e*_ in the insula has disappeared.

The distribution of each parameter across the cortical ribbon voxels is plotted in Figure 2.A. We note a bimodality of NEXI_NPA_ for the estimation of fraction *f*, but also more estimates close to the upper bounds. Thus, the main modes of *t*_*ex*_, *f* and *D*_*e*_ are located at lower values than the range displayed for the brain surfaces (Fig.1) for NEXI_NPA_, due to smoothing across this bi-modal distribution. Notably, the *t*_*ex*_ mode is located around 2 ms for NEXI_NPA_ and 11 ms for NEXI_WP_, the latter being closer to the values reported in human cortex to date using Connectom scanners and shorter gradient pulses (Chan et al., 2024; Uhl et al., 2024b).

**Fig. 2.**
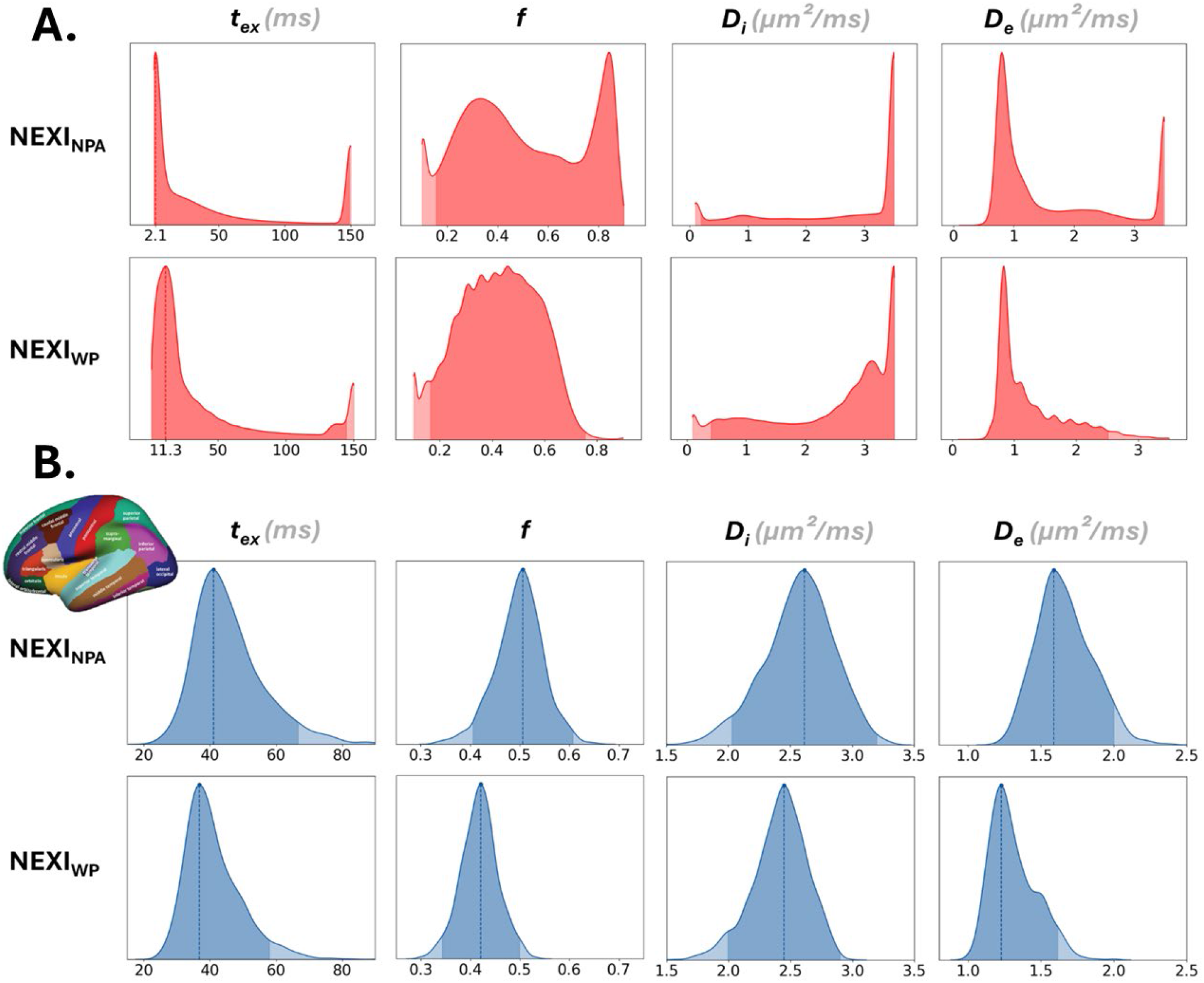
**A**. Parametric distributions across voxels in the cortical ribbons from NEXI_NPA_ and NEXI_WP_ estimations. The NEXI_NPA_ model most often gives values at the boundaries and produces a bimodality in *f*, which completely disappears with the other NEXI variant. The NEXI_WP_ fit shows comb-like peaks that result from the combination of the ODE solver’s imprecision with the initial grid-search. **B**. Distributions of ROI means from the DKT atlas (top-left illustration), from NEXI_NPA_ (top row) and NEXI_WP_ (bottom row) estimations. The values obtained with NEXI_WP_ are lower than those with NEXI_NPA_.

To gain insight at the ROI-level, we also examine the NEXI parameter distributions across the 68 ROIs of the DKT atlas averaged within each ROI and over all subjects (Figure 2.B). These distributions yield well-defined modes, which we report in Table 1, with the 95% confidence intervals obtained by integrating around the mode. The ROI-averaged values are in better agreement with the smoothed brain surface values in Figure 1.

**Table 1.**
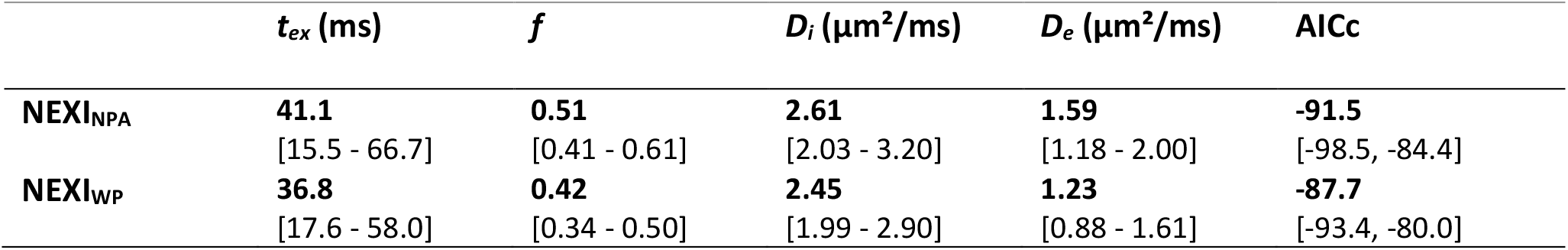
Modes and confidence intervals of NEXI estimates across DKT ROIs. The mode and confidence intervals of the corrected Akaike Information Criterion (AICc) are also given.

Finally, Table 1 also reports the mode and confidence intervals of the AICc of the two models. The full distribution of AICc values across the DKT ROIs is shown in Figure S3. A Wilcoxon signed-rank test was performed to compare the distributions of scores, which showed that NEXI_NPA_ AICc is significantly lower than NEXI_WP_ AICc, p < 0.0001. These results suggest that the signals derived from NEXI_NPA_ estimates are surprisingly closer to the experimental signals.

### 3.2 Fitting performance evaluation

#### 3.2.1 NEXI noise propagation: fitting synthetic ground truth signals

Estimation performance on the synthetic dataset is shown in Figure 3. As the underlying ground truth is generated considering the actual diffusion gradient pulse width, the accuracy of NEXI_WP_ estimates is good, while NEXI_NPA_ estimates are naturally more biased. For both variants, the accuracy on *t*_*ex*_ deteriorates with longer times, which is due to the limited diffusion time range sampled, and the precision is poorest on *t*_*ex*_ and *D*_*i*_, as previously reported (Jelescu et al., 2022). NEXI_NPA_ biases reflect differences in the distributions obtained on the experimental data. For example, *t*_*ex*_ values are underestimated while *D*_*i*_ is overestimated. *D*_*e*_ estimates are relatively unbiased in the vicinity of 1 µm^2^/ms, echoing the almost identical estimation modes of these two models on our experimental data. For *f* < 0.5, NEXI_NPA_ outputs are overestimated, consistent with the higher mode for *f* produced by this variant in the experimental data. Under equivalent data and fit, with δ reduced to 4 ms, NEXI_WP_ surprisingly exhibits greater bias than NEXI_NPA_, which has significantly diminished its own bias, as shown in Figure S7.

**Fig. 3.**
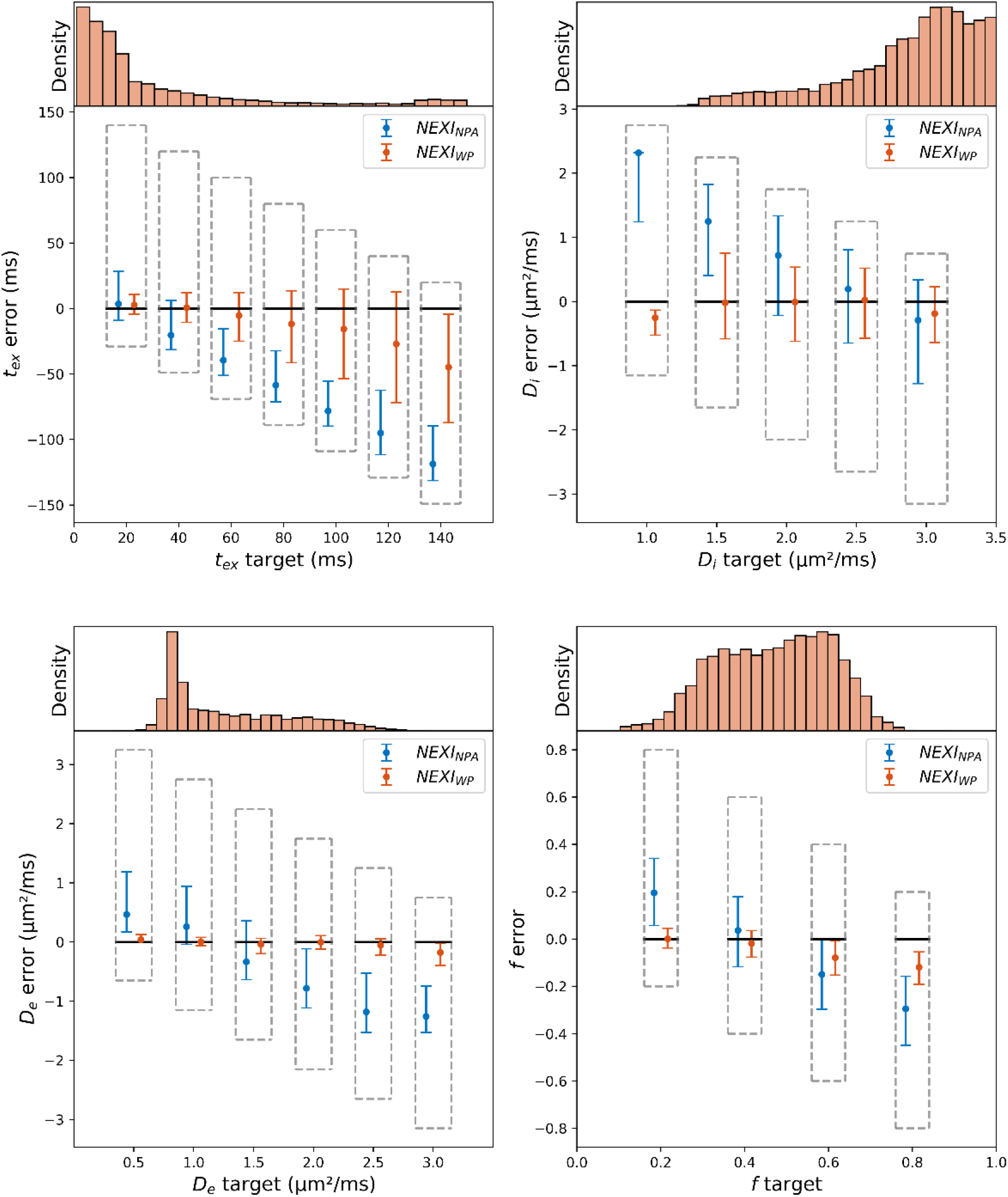
Boxplot (median and interquartile range) of NEXI_NPA_ and NEXI_WP_ parameter estimates from synthetic NEXI_WP_ signals generated using the experimental estimates from NEXI_WP_ as ground truth, and experimental Rician noise levels. The error is defined as the difference between the estimation and the target value. The upper and lower limits of the grey dashed box represent the range of possible errors allowed by the NLS bounds. *t*_*ex*_ and *D*_*i*_ exhibited the greatest variance, especially for NEXI_WP_. NEXI_NPA_ showed more bias, systematically underestimating *t*_*ex*_ and overestimating *D*_*i*_.

#### 3.2.2 Reproducibility

To assess intra-subject variability, we compared the first and second sessions of all our subjects for each DKT ROI and plotted the distribution of these differences. To assess inter-subject variability, we compared the first session of each subject with those of the others. We show these distributions for *t*_*ex*_ and *f* (Figure 4A) and for the diffusivities (Figure 4B) from both NEXI implementations. A Wilcoxon signed-rank test showed that the differences between scans but also between subjects are smaller for NEXI_WP_ than NEXI_NPA_ (p < 0.0001). NEXI_WP_ also improves intra-subject correlations compared to NEXI_NPA_, as show in Table 2. Both models however display modes and means of the estimation difference distributions that are greater for inter-subject differences than intra-subject differences indicating that the models are sensitive to inter-subject variations. The inter-subject and inter-session difference can also be seen in the less pronounced inter-subject correlations presented in Table 2.

**Table 2.**
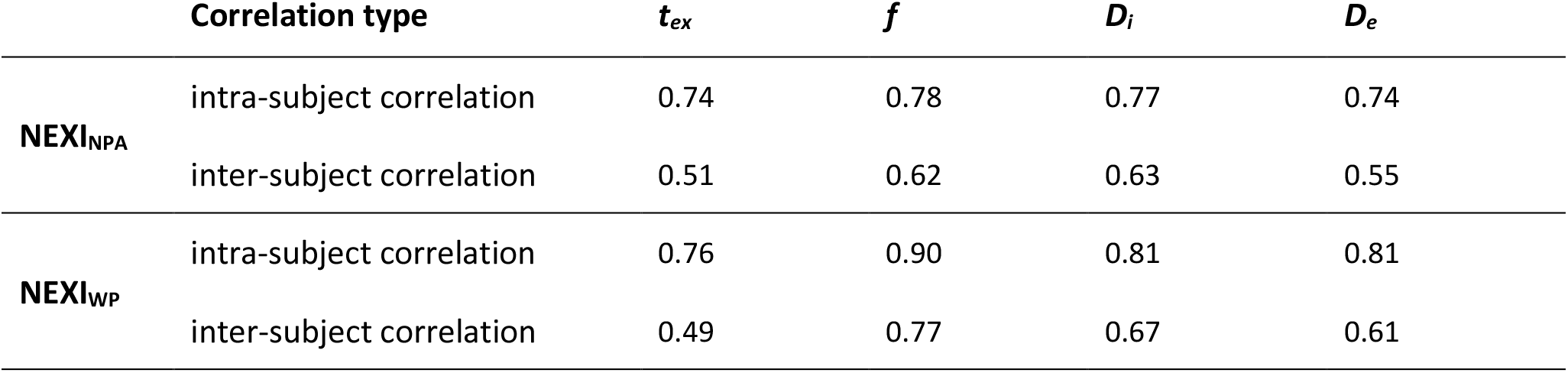
Correlation coefficients of the DKT ROI means between the different sessions of all subjects (denoted intra-subject) and between the different subjects first session (denoted inter-subject). The correlation is less pronounced between subjects than for several sessions of the same subject. The NEXI_WP_ model also shows stronger correlations both intra and inter subjects, except for *t*_*ex*_. All correlations verified p<0.0001.

**Fig. 4.**
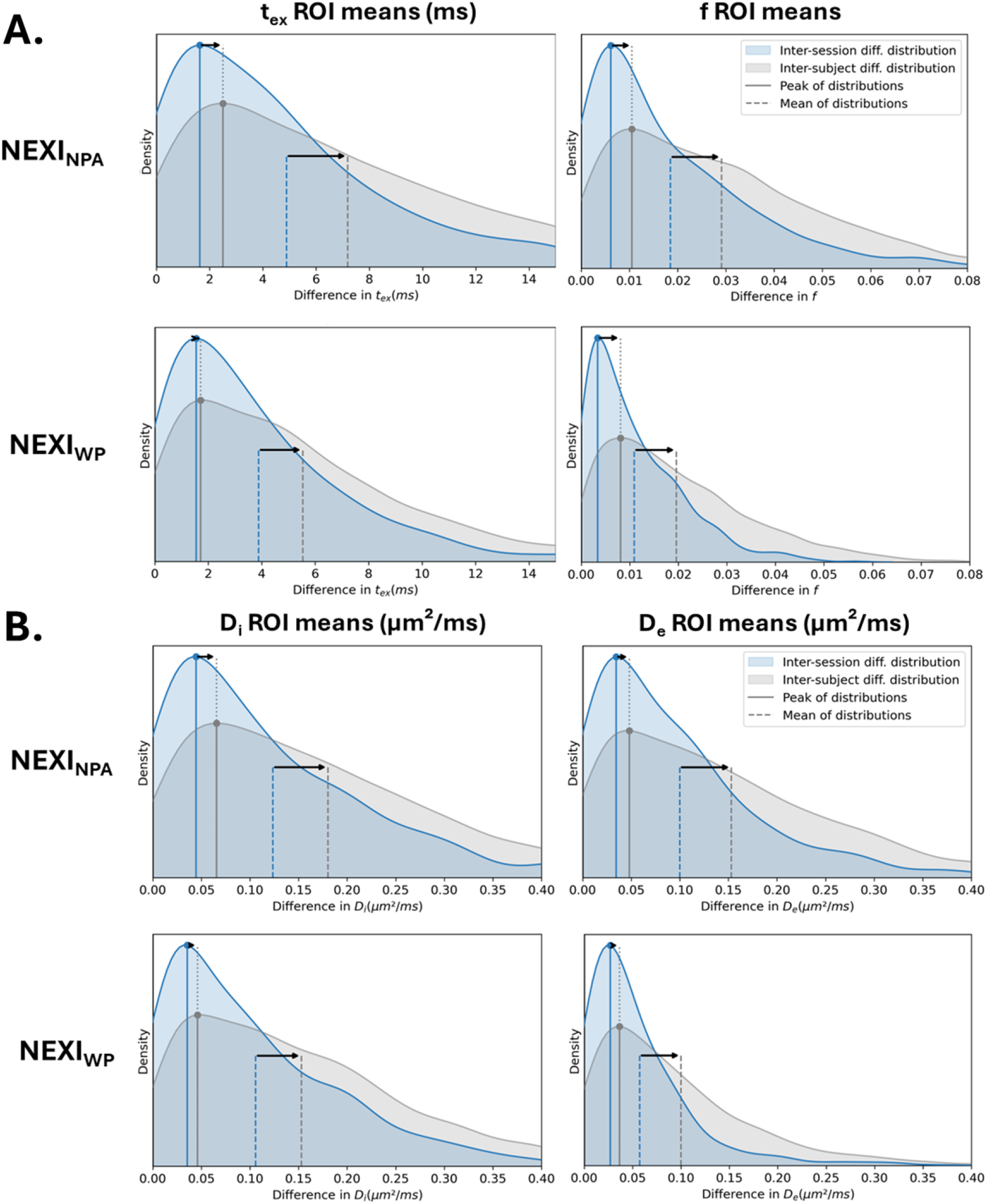
**A**. Distributions of the absolute differences of the DKT ROI means of NEXI_NPA_ and NEXI_WP_ *t*_*ex*_ and *f* estimates between two sessions of the same subjects (blue) and between two subjects (gray). **B**. Equivalent distributions for *D*_*i*_ and *D*_*e*_ estimates. The modes/peaks seem to vary less between inter- and intra-subject distributions, the difference being more in the tail of the distribution, which shifts the overall mean of the differences.

### 3.3 Correspondence between NEXI parameters and the MWF

The correlations between each NEXI parameter and the MWF across DKT ROIs are shown in Figure 5. The exchange time shows a significant correlation to MWF (r=0.74, 0.76, respectively for NEXI_NPA_ and NEXI_WP_), followed by the extra-cellular diffusivity *D*_*e*_ (r=0.41, 0.55), and to a lesser extent *f* (r=−0.37, −0.30) and *D*_*i*_ (r=−0.31, −0.36). The correlation coefficients and p-values are very similar between the two model variants, showing the same trends.

**Fig. 5.**
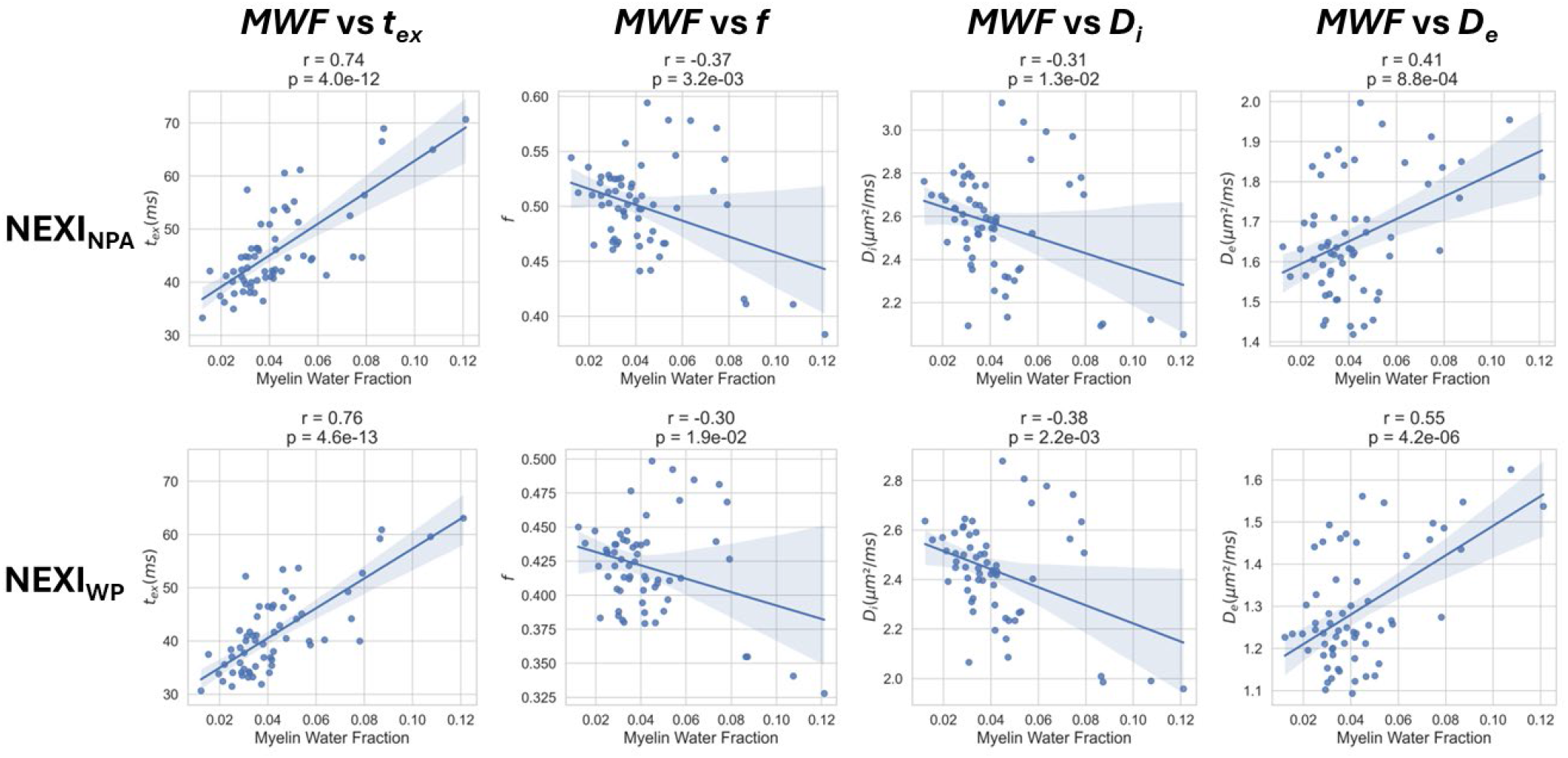
Correlation between average MWF of the DKT ROI and average parameters of NEXI_NPA_ and NEXI_WP_. In both NEXI models, there is a very significant strong positive correlation between MWF and *t*_*ex*_, a moderate positive correlation with *D*_*e*_, and weak negative correlations with *f* and *D*_*i*_.

The strong relationship between *t*_ex_ and MWF is also reflected in the patterns observed on cortical surfaces projections. MWF and *t*_ex_ projections are plotted in Figure 6 where we provide the *T*_*1*_*w/T*_*2*_*w-ratio* Myelin Fraction map from (Glasser et al., 2016) as reference for comparison. Similar patterns of elevated MWF and longer *t*_ex_ can be observed in the primary motor cortex and other areas proximal to the central sulcus. However, the higher *t*_*ex*_ patterns in the anterior temporal lobe are not matched in the MWF map, nor in the reference Myelin Map. This indicates that, despite the strong correlation, the MWF is not sufficient to explain the variations in *t*_*ex*_ across the cortex. Overall, the MWF map obtained through T_2_ relaxometry and the Myelin Maps show some consistent patterns mostly in the areas round the central sulcus, however, the Myelin map patterns found in the parietal, occipital (V1, MT) and the temporal lobe (A1) are not present in the MWF.

**Fig. 6.**
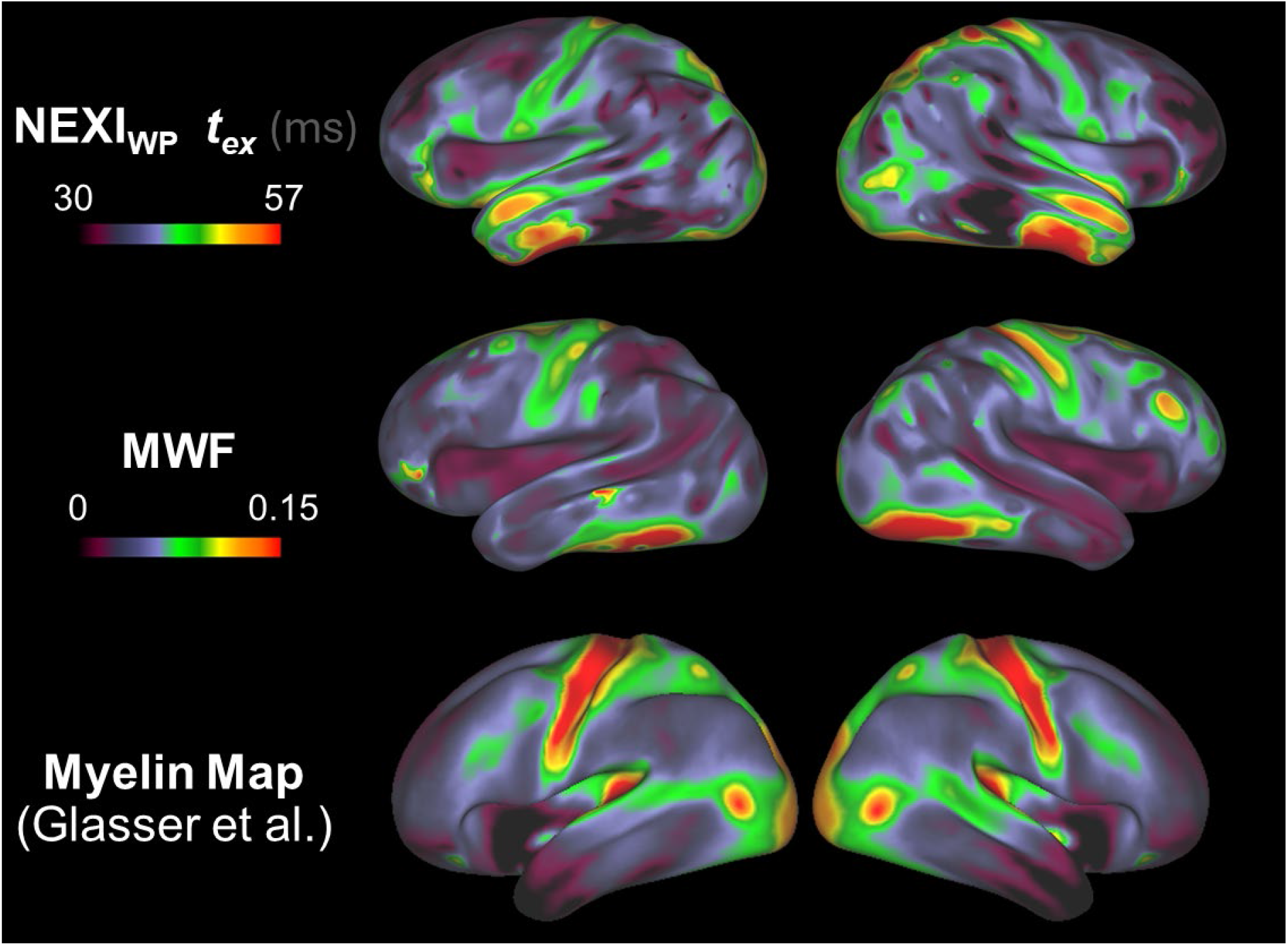
Projection onto cortical surface of NEXI_WP_ *t*_*ex*_ estimations and the MWF. A reminder of the Cortical Myelin Maps using T_1_w/T_2_w ratio obtained in (Glasser et al., 2016) is shown below for reference. Both the *t*_*ex*_ and MWF maps show higher values in the motor cortex and along the central sulcus, which is also largely present on the Myelin Map. The maps show differences in the temporal lobe.

### 3.3 Correspondence with cortical thickness

The correlations between NEXI parameters and cortical thickness across ROIs is shown in Figure S4, as well as a comparison of cortical surface maps in Figure S5. The intra-neurite fraction shows a significant correlation to cortical thickness (r=−0.74, −0.78, respectively for NEXI_NPA_ and NEXI_WP_), closely followed by the intra-cellular diffusivity *D*_*i*_ (r=−0.68, −0.66), and to a lesser extent the extra-cellular diffusivity *D*_*e*_ (r=−0.45, −0.36). Remarkably, *t*_ex_ does not correlate with cortical thickness. The patterns of *f* and cortical thickness across the brain surface confirm these two measures to be strongly anti-correlated.

## Discussion

In this study, we present the first in vivo quantification of human cortical microstructure using the NEXI model on data acquired with a clinical MRI scanner. In our previous NEXI implementation on Connectom data (Uhl et al., 2024b), we established that the model estimates greatly benefitted from accounting for the Rician mean floor in the fitting procedure. One of the open questions of our previous work was how NEXI could translate to clinical systems, given that weaker gradient strength and different stimulation limits needs to be compensated by longer gradient pulse durations to achieve high b-values with short diffusion times. Here we therefore compared two implementations of NEXI: NEXI_NPA_ (Jelescu et al., 2022), which approximates diffusion gradients as instantaneous pulses, and NEXI_WP_ (Olesen et al., 2022), which integrates the actual gradient waveforms for greater accuracy at the cost of a greater computation time. Both implementations already account for the Rician floor. The comparison was based on experimental data in the human cortex, as well as noise propagation in synthetic data. As in (Uhl et al., 2024b), we also examined intra-vs inter-subject variability of NEXI estimates. The sensitivity to individual differences is an important feature for clinical translation. Finally, we add to the validation of NEXI estimates against complementary measures by examining the correlations to MWF and cortical thickness.

Our findings reveal that both NEXI_NPA_ and NEXI_WP_ yield parameter estimates consistent with previous studies on human gray matter using the Connectom scanner (Uhl et al., 2024b) and in vivo rat cortex (Jelescu et al., 2022). Cortical features previously quantified on the Connectom scanner were, on average: *t*_*ex*_ = 42 ms, *f* = 0.38, *D*_*i*_ = 3.35 µm^2^/ms, *D*_*e*_ = 0.92 µm^2^/ms, which is largely similar to values reported here in Table 1. Notably, *t*_*ex*_ consistently falls within the range of tens of milliseconds.

While the overall spatial patterns of estimated NEXI parameters across the brain remain consistent between the two variants, NEXI_WP_, with its more precise gradient integration, tends to produce lower mean parameter values compared to NEXI_NPA_, and largely resolves the bimodality of solutions across all cortical voxels, particularly for *f* and *D*_e_. Although the AICc of NEXI_NPA_ is somewhat lower, i.e. the fit of this model described the signal better, NEXI_WP_ results seem more plausible from a physical point of view. For example, the distribution of *D*_i_ values across all cortical voxels displays fewer extreme values (higher than the free water diffusivity at 37°C), resulting in a mean *D*_i_ value over DKT ROIs of 2.45 µm^2^/ms, which is similar to axonal diffusivity reported in white matter (Dhital et al., 2019b; Howard et al., 2022b; Tristão Pereira et al., 2021).

Using synthetic data, we examined how the NEXI_NPA_ model estimates are impacted by long pulses. For most parameters, notably *D*_*i*_, *D*_*e*_ in the low ranges and *f* in the low ranges, the simulation reproduces the differences observed experimentally: NEXI_NPA_ estimates are higher than NEXI_WP_. However, while in data the exchange time estimate is longer for NEXI_NPA_ than NEXI_WP_, the simulation suggests that NEXI_NPA_ underestimates *t*_ex_ in the case of a wide pulse ground truth. We do not have an explanation for these divergent observations. These results are not observed when δ is reduced to 4 ms. In this scenario, NEXI_NPA_ significantly reduces the bias in its estimates, while NEXI_WP_ maintains biases and variances similar to those seen with longer δ. This suggests that NEXI_WP_ bias may stem from the estimator itself, whereas NEXI_NPA_ bias likely arises from the violation of the short-pulse assumption.

The differences in inter-subject and intra-subject correlations suggest that both models can be sensitive to inter-subject differences. With NEXI_WP_, the distinction between inter- and intra-subject differences is less pronounced. While intra-subject correlations are expected to be as high as possible (and they are higher with NEXI_WP_ than with NEXI_NPA_), expected inter-subject correlations are difficult to estimate. It is therefore possible that one or other of the inter-subject correlations between NEXI_NPA_ and NEXI_WP_ is the most accurate.

Surprisingly, NEXI_NPA_ has lower AICc and therefore better fitting. However, a better fit does not always guarantee a better model, especially as the AICc confidence intervals of the two variants overlap.

The discrepancy between NEXI_NPA_ and NEXI_WP_ estimates, particularly in the exchange time, raises questions about the validity of the narrow pulse approximation in clinical settings with longer gradient durations. While NEXI_NPA_ offers a computationally efficient approach for parameter estimation, its accuracy may be compromised as shown by the simulation. NEXI_WP_, on the other hand, provides a more accurate representation of the diffusion process but at the cost of increased computational complexity. The choice between NEXI_NPA_ and NEXI_WP_ may depend on the specific research question and the available computational resources. The use of NEXI_WP_ may be particularly important in studies investigating subtle changes in gray matter microstructure, such as those associated with early stages of neurodegenerative diseases, where accurate parameter estimation is crucial. It is probably just as crucial to take the pulse duration into account when large cohorts of patients with different protocols with different δ are analyzed.

Our analysis reveals a significant positive correlation between *t*_*ex*_ derived from both NEXI versions and the MWF, characterized by a very high correlation coefficient of 0.7. The exchange time is the NEXI parameter that correlates most strongly with the MWF. This correlation aligns with the expectation that regions with higher myelin content exhibit longer exchange times due to reduced membrane permeability. The presence of myelin acts as a barrier to water diffusion, hindering the exchange of water molecules between the intra- and extra-neurite compartments. This is supported by previous studies that have shown a relationship between myelin content and restricted diffusion in white matter (Dhital et al., 2019b; Lampinen et al., 2020).

Spatially, *t*_*ex*_ maps indeed displayed patterns of slower exchange where the Myelin Map showed increased myelin density (Glasser et al., 2016). These patterns were localized around the central sulcus (primarily motor M1 but also somatosensory S1) and in other primary areas such as visual V1 and auditory A1, known for their higher myelin content due to large sensory projections (Glasser et al., 2014; Nieuwenhuys and Broere, 2017), and high-functioning visual areas like middle temporal visual area MT (Born and Bradley, 2005; Glasser et al., 2014; Sánchez-Panchuelo et al., 2012; Sereno et al., 2013).

On the other hand, the anterior temporal lobe showed slower exchange with NEXI_WP_ while the MWF map showed lower myelin density, similarly to cytoarchitectonic studies also reporting lower myelin content in the area (Glasser et al., 2014; Shafee et al., 2015). This suggests that factors beyond myelination also contribute to variations in exchange time.

The high *t*_ex_ values found in the temporal areas are surprising, especially as they do not seem to be explained by myelin content nor partial volume with subcortical white matter (as the cortex is in fact particularly thick in the temporal lobe). The correlation between *t*_*ex*_ and MWF could also be influenced by the density of other cell types in the cortex, such as astrocytes, whose processes likely contribute to the “neurite” compartment. Indeed, NEXI maps in the rat brain that were compared against immunohistochemistry staining had revealed that neurite density patterns showed a good correspondence with neurofilament NeuF staining across cortical layers, but rather with astrocyte GFAP staining in hippocampal sub-fields (Jelescu et al., 2022). The density of dendritic spines may also influence the estimate of exchange time across the cell membrane, as the spines may act as a different environment that is in “exchange” with the core of the dendrite (Chakwizira et al., 2024; Şimşek and Palombo, 2024).

The central sulcus, characterized by both high *f, D*_*i*_ and low cortical thickness, exemplifies the potential influence of partial volume effects on NEXI parameter estimation. The thinner cortex in this region may lead to increased partial volume with white matter, which could affect the estimates. Since WM is characterized by long impermeable myelinated fibers, the presence of white matter in the same voxel as gray matter could lead to higher estimates of *t*_*ex*_, *f* (and possibly *D*_*i*_), as well as MWF. Remarkably, across the whole cortex, the DKT means of *f* and *D*_*i*_ strongly correlate with the cortical thickness, but not *t*_ex_ nor MWF. This indicates once again that PVEs, and hence the contribution of WM and therefore myelin to the voxel, are not sufficient to explain changes in *t*_*ex*_. The significant correlation between cortical thickness and NEXI parameters could also be investigated using a surface-based analysis, which may be more sensitive to local variations than a region-of-interest-based analysis.

In addition to the factors mentioned above, the estimation of NEXI parameters may also be influenced by other microstructural features not explicitly modeled by NEXI, such as the presence of non-Gaussian diffusion within compartments due to irregularities like dendritic spines and neurite beading (Henriques et al., 2019; Lee et al., 2020). Future studies should investigate the impact of these features on NEXI parameter estimation and explore the potential of incorporating them into the model to improve its accuracy and specificity. Such a model has been developed recently (Novikov et al., 2023) but has not yet been applied to clinical nor preclinical data. The inclusion of a soma compartment, as proposed in the SANDIX model (Olesen et al., 2022) or GEM model (Uhl et al., 2024a), could also be explored to account for the contribution of cell bodies to the diffusion signal in gray matter. However, the inclusion of additional compartments will increase the complexity of the model and make it more challenging to estimate its parameters with sufficient accuracy and precision. The choice of the appropriate model complexity should be guided by the specific research question and the available data quality.

Furthermore, the high values of *D*_*i*_ observed in our study, at times exceeding the water diffusion coefficient at body temperature, suggest that the model may be overestimating this parameter. This finding was already noted in our previous Connectom study, although the NEXI_WP_ model tends to mitigate it (Uhl et al., 2024b). This parameter is notoriously difficult to estimate for all models (Howard et al., 2022b; Jelescu et al., 2016a; Palombo et al., 2020). This could be due to several factors, including the presence of noise, partial volume effects, and the limitations of the model in capturing the complex diffusion environment within neurites. Future studies should investigate these factors in more detail and explore methods to improve the accuracy of *D*_*i*_ estimation, such as trying more realistic models of neurite geometry and diffusion. The assumption of uniaxial diffusion within neurites may not hold in regions with complex neurite architectures, due to the presence of dendritic spines and other neurite irregularities that are not captured by the current NEXI model.

The comparison of NEXI parameter estimates with histological data is crucial for validating the model and interpreting its parameters in terms of underlying tissue microstructure. However, such comparisons are challenging due to the limited availability of high-resolution histological data in the human brain. Future studies will aim to acquire high-quality histological data in conjunction with NEXI measurements on postmortem human brains to establish a more direct link between model parameters and tissue microstructure (Hertanu et al., 2023).

Another crucial factor in the adoption of the NEXI model is the time required to obtain reliable results. A short protocol is more likely to be integrated into a clinical routine, enabling the acquisition of data from patients presenting neurological alterations that this model could detect. Our next step will therefore be to reduce this protocol while maintaining good performance of the fit (Uhl et al., 2023).

## Conclusion

This study is the first to demonstrate the feasibility of estimating NEXI parameters in the human cortex in vivo using a clinical MRI scanner. Our findings reveal that NEXI_NPA_ and NEXI_WP_ provide similar gray matter microstructure parametric maps, with NEXI_WP_ offering potentially more accurate and biologically plausible results due to its consideration of wide diffusion pulses. The strong correlation between the exchange time and MWF further supports the biological relevance of NEXI parameters and the potential of t_ex_ as biomarker for cell membrane permeability – whether related to myelination or cell integrity – in the brain.

Future work will focus on shortening acquisition protocols for clinical scanners and exploring the potential of NEXI as a biomarker for various neurological conditions. The development of stronger gradients on clinical scanners and advancements in denoising techniques are expected to further improve the accuracy and precision of NEXI estimates, paving the way for its wider clinical application.

## Data and code availability

The code used in this study is available on https://github.com/Mic-map/greymatter_swissknife. The code for the Myelin Water Fraction estimation (Canales-Rodríguez et al., 2021) used in this study is available at https://github.com/ejcanalesr/multicomponent-T2-toolbox. The data used in this study are available upon request after signing a formal data sharing agreement and providing approval from the requesting researcher’s local ethics committee.

## Author contributions

Conceptualization: IJ, QU; Data curation: QU; Formal analysis: QU; Funding acquisition: IJ; Investigation: QU; Methodology: QU, TP, TF, GFP, IJ; Supervision: IJ; Visualization: QU, TP; Writing—original draft: QU; Writing—review & editing: QU, TP, TF, GFP, IJ.

## Acknowledgments

The authors thank Sune Jespersen for insightful discussions and for sharing the initial code for wide pulses. We acknowledge the CIBM Center for Biomedical Imaging for providing resources to conduct this study.

