## Supplementary material for "Human gray matter microstructure mapped using Neurite Exchange Imaging (NEXI) on a clinical scanner"

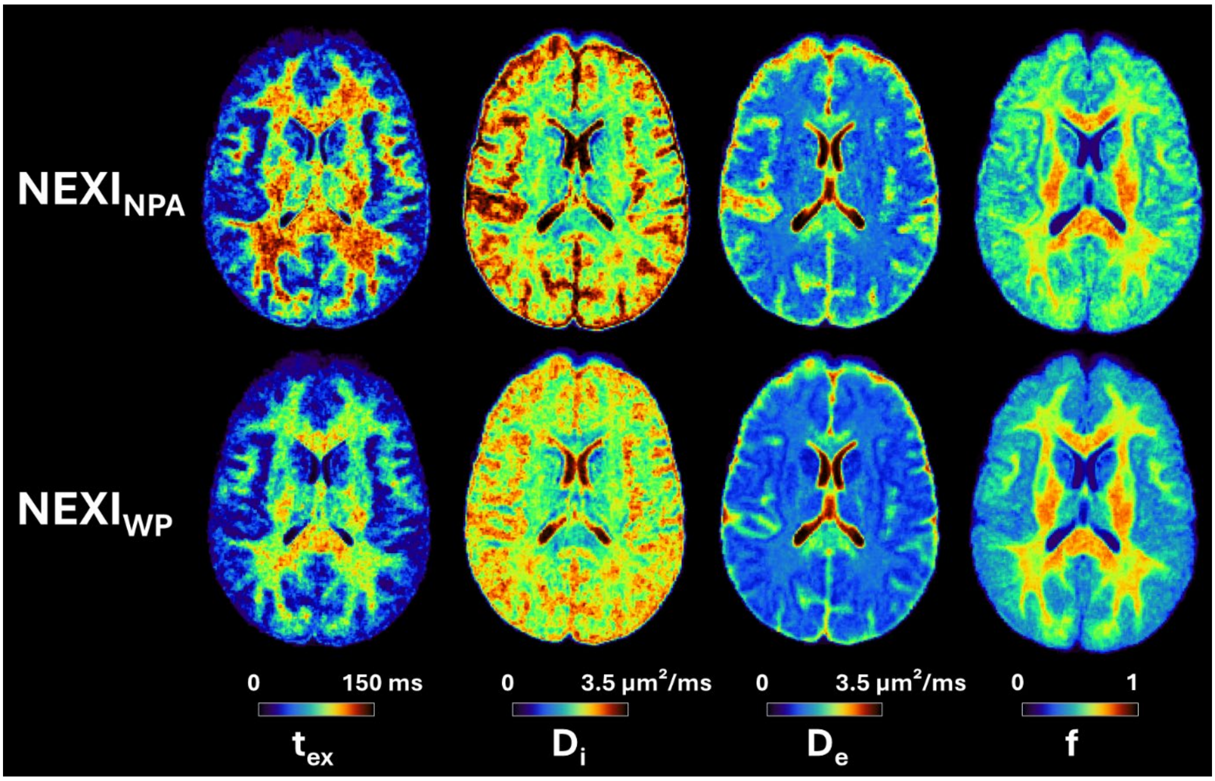


**Fig. S1** Axial slice of $NEXI_{NPA}$ and $NEXI_{WP}$ parametric maps, averaged across sessions and subjects (N = 22). $t_{ex}$ and $D_e$ are consistent throughout the cortex, but $t_{ex}$ is presumably longer in the WM and cannot be reliably estimated using available diffusion times. $f$ and $t_{ex}$ display the expected anatomical pattern in white versus gray matter. $D_i$ shows large variability across voxels, while hitting its upper bound frequently.

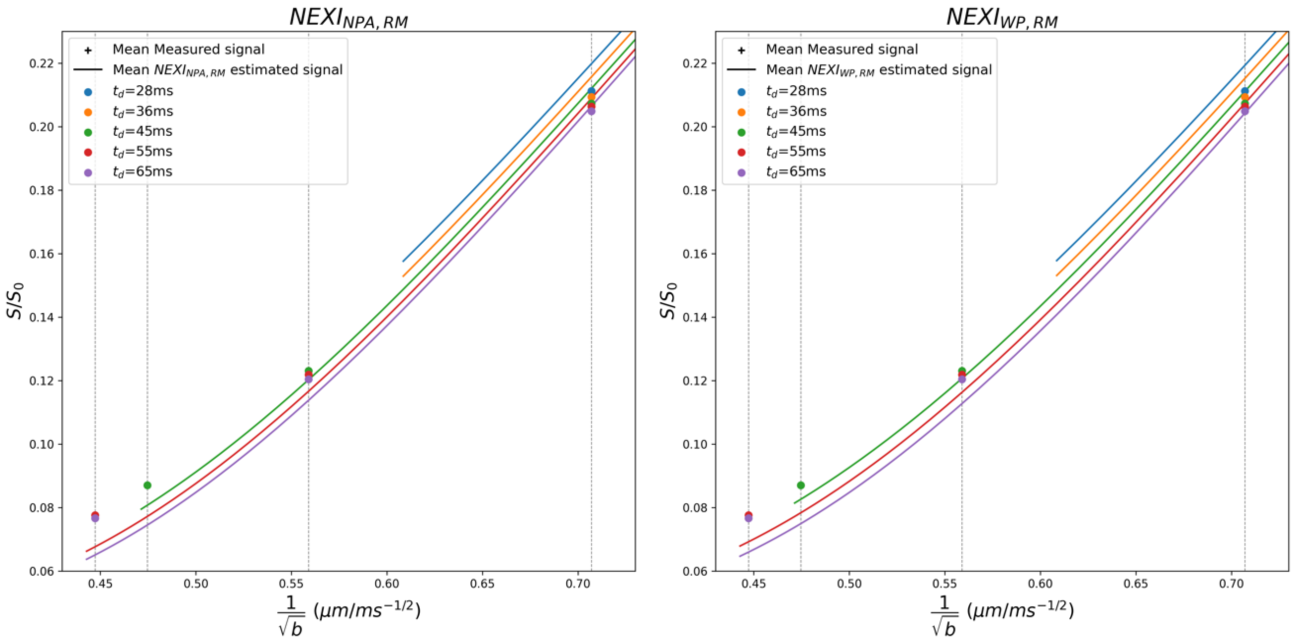


**Fig. S2.** Mean measured signal, represented by dots, and mean estimated signal curves of the whole cortical ribbon from both NEXI implementations. High b-value signals are better estimated with $NEXI_{WP}$. At low $t_d$, the measured signals have lower maximum b-value. Thus, at b=5 ms/µm², on the left on the x-axis, we only have signals from $t_d$ at 55 and 65 ms, at b=4.44 ms/µm², only 45 ms and at b=3.2 ms/µm² we have $t_d$ at 45, 55 and 65 ms. The low $t_d$ curves are therefore only partially shown.

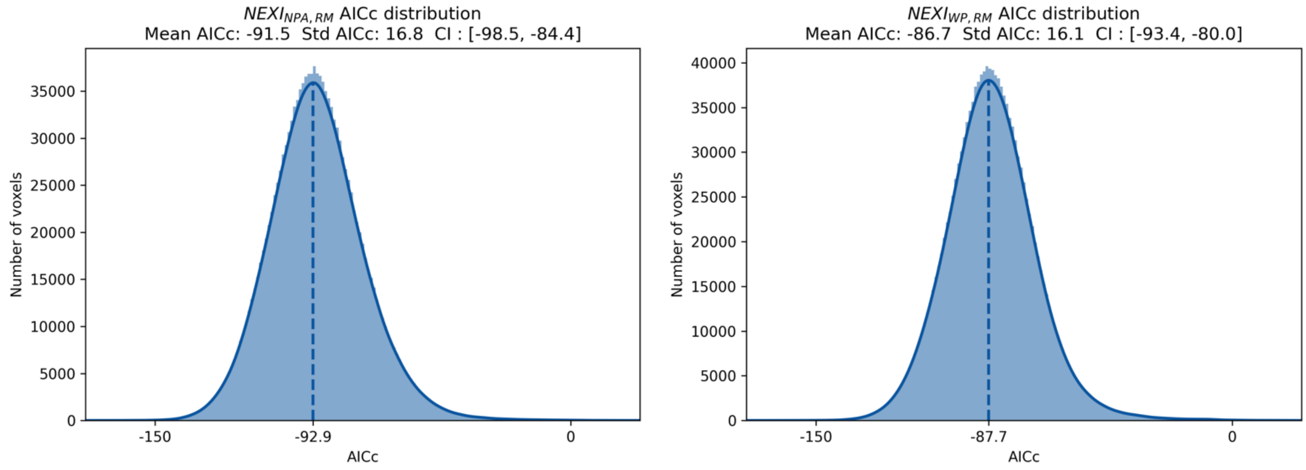


**Fig. S3.** Corrected Akaike Information Criterion (AICc) distribution of $NEXI_{NPA}$ and $NEXI_{WP}$ in the whole cortical ribbon. The performance of $NEXI_{NPA}$ and $NEXI_{NPA}$ is comparable, and their confidence intervals overlap.

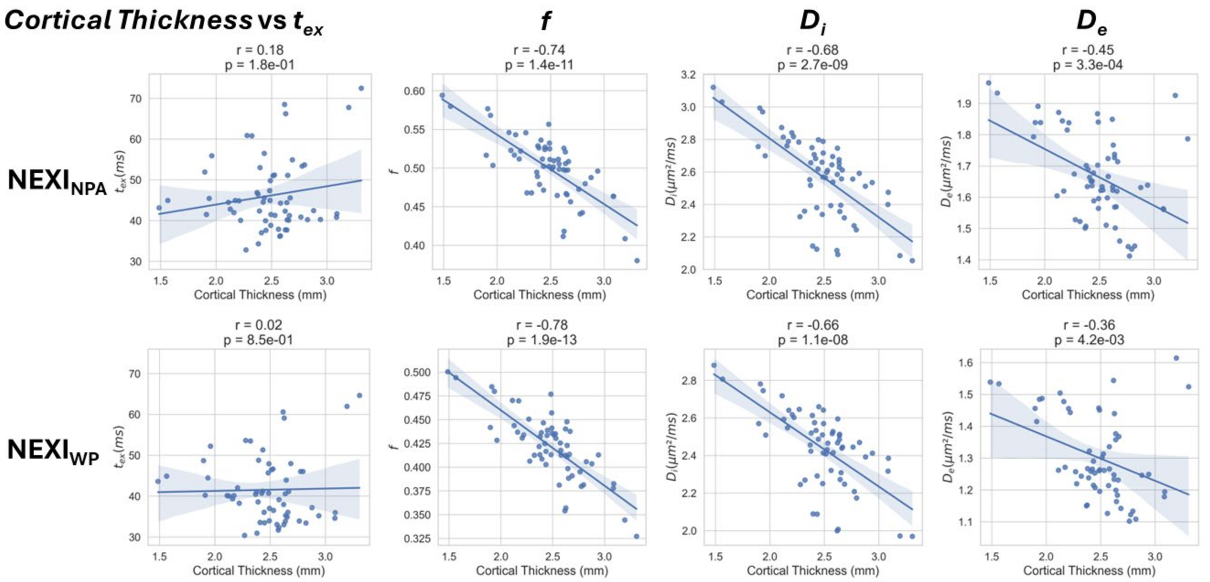


**Fig. S4** Correlation between DKT ROI means of Cortical Thickness and NEXI parameters in both $NEXI_{NPA}$ and $NEXI_{WP}$ implementations. There is a very significant and strong negative correlation between cortical thickness and f as well as with $D_i$.

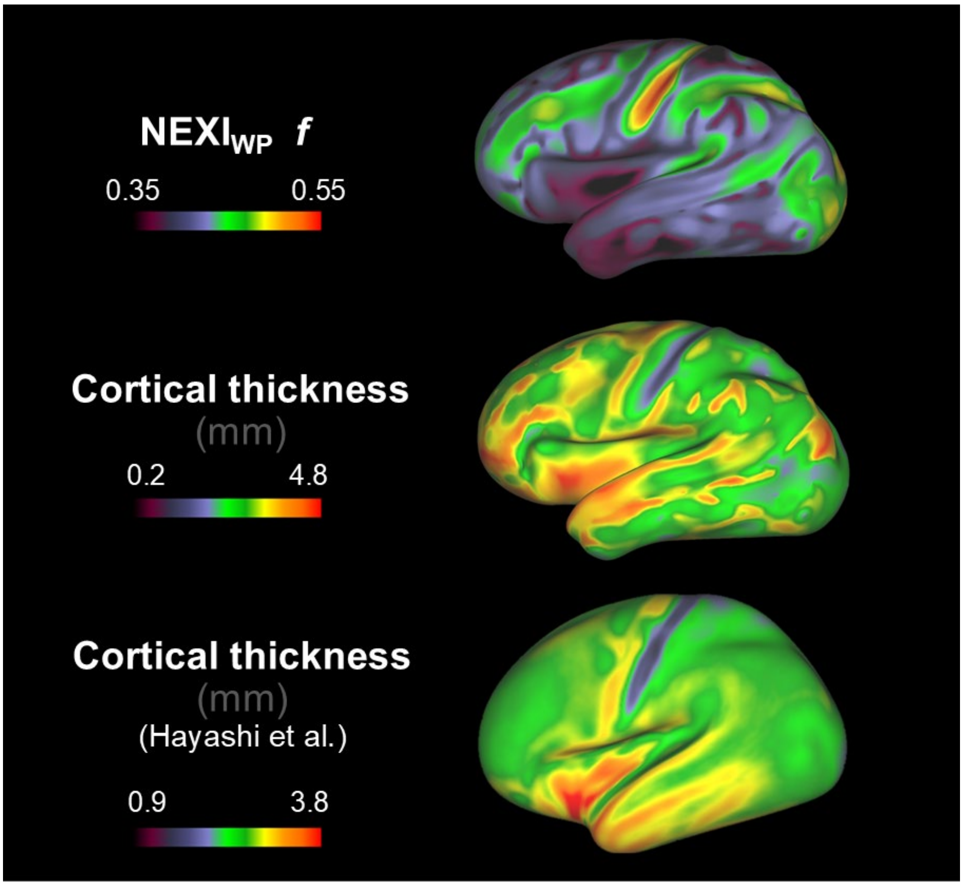


**Fig. S5** Projection onto cortical surface of $NEXI_{WP}$ *f* estimations and the cortical thickness. A reminder of the cortical thickness maps obtained in (Hayashi et al., 2021) is shown below for reference. The maps show similar trends. Especially, the main decrease in cortical thickness around the central sulcus is matching with the main increase in *f* in this region.

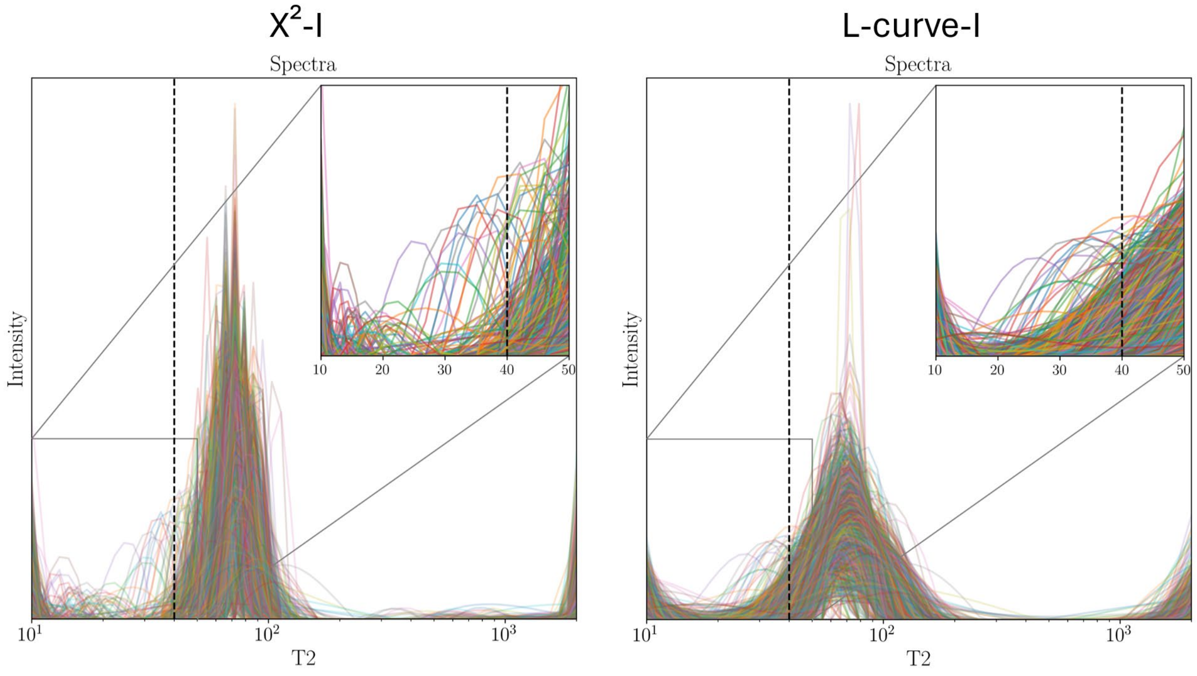


**Fig. S6** Comparison of the MWF cutoff of the intensity $T_2$ spectrum lobes from one subject from $\chi^2$-I and L-curve-I methods. These methods are recommended for MWF extraction respectively for moderate and high level of noise. The lobe separation of the $T_2$ spectrum reveals that the $\chi^2$-I method is better suited to our data.

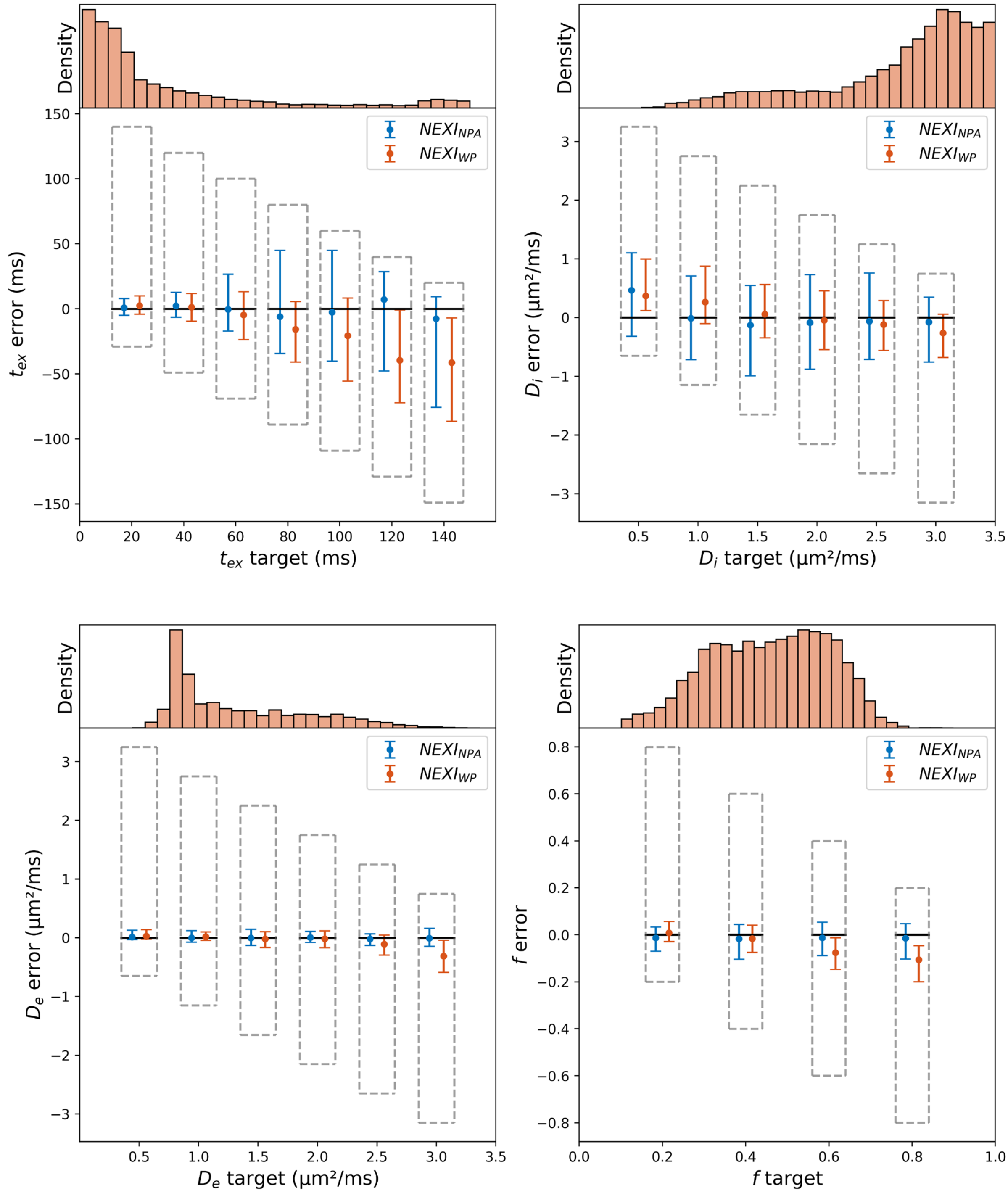


**Fig. S7** Boxplot (median and interquartile range) of $NEXI_{NPA}$ and $NEXI_{WP}$ parameter estimates from synthetic $NEXI_{WP}$ signals generated using the experimental estimates from $NEXI_{WP}$ as ground truth, and experimental Rician noise levels, as in Fig. 3, but setting δ to 4 ms. This time, $NEXI_{WP}$ showed more bias than $NEXI_{NPA}$.
